# Differential Effects of Cocaine Self-Administration Regimens on Incubation of Cocaine Craving and Nucleus Accumbens Neuronal Ensembles Activated by Cocaine-Associated Context

**DOI:** 10.1101/2025.03.17.643658

**Authors:** Augusto Anesio, Giovanna Victoria Costa Lopes, Paola Palombo, Caroline Riberti Zaniboni, Thais Suemi Yokoyama, Vincent Vialou, Cássio Morais Loss, Fábio Cardoso Cruz

**Author notes:** These authors contributed equally to this work (Cássio Morais Loss, Fábio Cardoso Cruz). Corresponding author: Fábio Cardoso Cruz.

## Abstract

Drug addiction develops in a subset of users following repeated exposure, influenced by biopsychosocial factors. Rodent self-administration protocols, varying in drug access times, are used to study both controlled and compulsive drug-taking behaviors and their neurobiological underpinnings. Drug-associated cues and environmental contexts are well-established triggers for relapse, with susceptibility to these stimuli peaking during early abstinence and remaining elevated, thereby increasing relapse risk. This phenomenon, known as incubation of craving, has been replicated across substances in animal models. The associative learning between drug effects and contextual cues is encoded by neuronal ensembles activated by drug-associated stimuli, driving craving, seeking, and relapse. Neuronal ensembles in the nucleus accumbens (NAcc) are strongly involved in drug seeking, with parvalbumin-expressing fast-spiking interneurons playing a key role in this associative learning process. We investigated activation patterns in the NAcc triggered by cocaine-related cues following restricted- or extended-access self-administration and examined how forced abstinence alters these patterns, contributing to the incubation of cocaine craving. We also analyzed the engagement of parvalbumin-positive interneurons in NAcc neuronal ensembles before and after forced abstinence. Our findings show that the extended access protocol more effectively induced the incubation of cocaine craving. Neuronal activation in the NAcc core increased after thirty days of forced abstinence in both groups, with extended access rats showing consistently higher activation. Forced abstinence also increased NAcc shell activation, with no differences between protocols. NAcc core activation, but not shell, was influenced by cocaine consumption during training. Notably, extended access rats exhibited reduced parvalbumin interneurons activation following thirty days of forced abstinence. Based on these findings, we speculate that the transition from occasional to compulsive drug-taking may be driven by molecular changes in the NAcc core that enhance its responsiveness to drug-related cues. Additionally, the incubation of craving could be linked to impaired inhibitory control of NAcc core medium spiny neurons by PV interneurons.

## Introduction

Drug addiction often develops after repeated drug use, marked by a gradual increase in consumption and a loss of control over drug intake, besides negative consequences of the use. Only a minority of individuals who repeatedly use drugs progress from occasional to compulsive drug taking. The transition from occasional, controlled drug use to addiction is thought to result from the interaction of biopsychosocial factors, and its neurobiological substrates are not yet fully understood (Ahmed & Koob, 1998; Deroche-Gamonet & Piazza, 2014; Kalivas & O’Brien, 2008; Thomas et al., 2009).

Despite their limitations in addressing the psychosocial aspects of addiction, animal models are valuable for understanding the neurobiological changes arising from the interaction between the individual and repeated drug use in the development of a compulsive phenotype (Field & Kersbergen, 2020; Spanagel, 2017). The self-administration paradigm is a wide-used model for investigating drug-taking behavior. It is based on the subject’s free choice and the influence of contextual cues on drug-seeking, allowing the exploration of the neurobiological changes arising from the interaction between the individual and repeated drug use in the development of a compulsive phenotype (Gardner, 2000; Panlilio & Goldberg, 2007).

By using distinct self-administration protocols, it is possible to investigate both controlled and compulsive drug-taking phenotypes, each associated with distinct molecular adaptations, shedding light on the potential mechanisms underlying drug addiction (Ahmed & Koob, 1998; Kawa et al., 2016; Wolf, 2016). Restricted-access protocols, with daily self-administration sessions lasting 1–2 hours, model behaviors consistent with occasional drug use. In contrast, extended-access protocols (4–6 hours daily) induce compulsive behaviors such as escalated drug consumption, persistent seeking without reinforcement, heightened motivation in progressive-ratio tests, disregard for danger cues, risk-taking, and increased relapse susceptibility triggered by priming or stress (Ahmed et al., 2000, 2002; Ahmed & Koob, 2004; Guillem et al., 2014).

Drug-associated cues and environmental contexts are well-established triggers for relapse (Cruz et al., 2013; Kalivas & O’Brien, 2008; Siegel, 1999; Wikler, 1973). Clinical studies reveal that the patients’ susceptibility to these stimuli increases during the initial weeks of abstinence and remains elevated for extended periods, contributing significantly to the high relapse rates observed within the first year of treatment (Dong et al., 2017; Gawin & Kleber, 1986; Pickens et al., 2011). This phenomenon, termed as incubation of craving, has been replicated in animal models across various substances, including cocaine, alcohol, nicotine, and methamphetamine (Abdolahi et al., 2010; Bedi et al., 2011; Bienkowski et al., 2004; Grimm et al., 2001; Li et al., 2015; Parvaz et al., 2016; G. Wang et al., 2013).

Crucially, the incubation of craving is characterized by heightened drug-seeking behavior following periods of forced abstinence (Abdolahi et al., 2010; Bedi et al., 2011; Bienkowski et al., 2004; Grimm et al., 2001; Li et al., 2015; Parvaz et al., 2016; G. Wang et al., 2013). Additionally, studies show that extended-access protocols effectively produce incubation of drug craving, whereas restricted-access protocols are less effective to produce such phenomenon (Wolf, 2016).

The aberrant behaviors seen in drug addiction are linked to long-lasting molecular changes in the brain, resulting in neuronal network reorganization. In this context, evidence shows alterations in the nucleus accumbens (NAcc), positioning it as a key region in the neuropathology of addiction due to its role as a central hub for integrating mesocorticolimbic signals that drive behavioral outcomes (Cornish & Kalivas, 2000; Engeli et al., 2021; Nestler, 2005; Wolf, 2010).

During drug use, dopamine may reinforce associations between specific environmental stimuli and the drug, enabling these cues to drive the behavior upon future exposure. The memory associations between drug-related effects and contextual cues are encoded by sparse, strongly interconnected groups of neurons, now called neuronal ensembles that become activated when the previously associated stimuli are encountered (Cruz et al., 2013; Di Chiara & Bassareo, 2007; Floresco, 2015; Guillem et al., 2014; Hebb, 1949; O’Donnell, 2003).

A growing body of evidence suggest that neuronal ensembles encoding addiction-related behaviors exhibit specific alterations compared to the general neuronal population. Moreover, selective manipulation of these ensembles effectively disrupts cue-induced drug-seeking behavior, which suggests that targeting specific neuronal ensembles, rather than entire brain regions, could provide valuable insights into the neurobiology of drug addiction and offer more precise therapeutic approaches (Cruz et al., 2013; Fanous et al., 2012; Koya et al., 2016; Xue et al., 2017).

Neuronal ensembles involved in encoding drug-associated memories within the nucleus accumbens are likely composed of its principal cell types, including medium spiny neurons (the predominant population in this region), alongside cholinergic and GABAergic interneurons. Among these GABAergic interneurons, parvalbumin-expressing (PV) fast-spiking interneurons play a key role in drug related associative learning. These neurons exert inhibitory control over medium spiny neurons in the nucleus accumbens, shaping neural dynamics essential for memory processing (Hijazi et al., 2023; Lee et al., 2017; X. Wang et al., 2018).

Accumulating evidence supports the involvement of PV interneurons in various aspects of memory. Specifically, during memory recall, PV interneurons help coordinate neural activity across limbic structures, including the nucleus accumbens, to facilitate the retrieval of appetitive memories and ensuring precise encoding and retrieval of memories (Hijazi et al., 2023; Trouche et al., 2019). Despite these findings, the specific role of PV interneurons in contextual cue-induced seeking behavior and incubation of craving remains poorly understood and requires further investigation.

The immediate early gene Fos expression is widely used to identify neuronal ensembles strongly activated in response to specific stimuli. Using this approach, we examined the activation patterns in the nucleus accumbens (NAcc) triggered by cocaine-related cues following restricted- or extended-access protocols. We further examined how forced abstinence alters these activation patterns to drive the incubation of cocaine craving, and to further understand circuit-level dynamics, we also analyzed the engagement of parvalbumin-positive interneurons in both protocols, before and after forced abstinence.

## Material and Methods

### Animals and environmental conditions

Adult male Wistar rats (postnatal day 60) were obtained from the Center for the Development of Experimental Models for Biology and Medicine (CEDEME) at the Federal University of São Paulo (UNIFESP) and were allowed to acclimate to the local vivarium for seven days before undergoing experimental procedures. They were group-housed (N = 4 per cage) and maintained under a 12:12-hour light-dark cycle (lights on at 6:30 AM) in a temperature-controlled environment (23 ± 2°C), with *ad libitum* access to food and water, except during self-administration sessions. The rats underwent surgery for intravenous catheterization and were single-housed until the end of the experiment to prevent catheter damage. Rats that maintained catheter patency until the experiment’s conclusion (N = 163 out of 189) and met the learning criteria during training (N = 120 out of 163) were included in the data analysis. Importantly, as some rats were allocated to other laboratory studies, only 71 out of the 120 included were used in the present study. The Research Ethics Committee of UNIFESP approved the experimental protocol (CEUA No. 4183030918).

### Drugs

Cocaine hydrochloride (95% purity) was provided by the Forensics Department of the São Paulo State Police (Process. 581/19 – DIPO). Cocaine hydrochloride was dissolved in sterile saline at individually adjusted concentrations for each animal, ensuring the delivery of 0.5 mg/kg per 0.1 mL infusion.

### Surgery for intravenous catheterization

Rats received preemptive analgesia 10 minutes before surgery with subcutaneous flunixin (2.5 mg/kg, 0.1 mL/100 g; Chemitec) and intraperitoneal fentanyl (0.05 mg/kg, 0.1 mL/100 g; Janssen). Anesthesia was induced via an intraperitoneal injection of a ketamine (80 mg/kg) and xylazine (10 mg/kg, 0.1 mL/100 g; Cristália) cocktail. Surgery commenced once the animals no longer exhibited reflex responses to paw pinching, which was repeatedly assessed throughout the procedure to ensure adequate anesthesia. Before making incisions, the surgical sites were shaved, cleaned, and disinfected with 70% alcohol and iodine. The catheter used for cocaine delivery (a silicone tube, internal diameter: 0.3 mm; external diameter: 0.64 mm; Dow Corning Corp) was inserted subcutaneously from the dorsal region toward the right jugular vein. A separate incision near the jugular vein exposed the vessel, allowing catheter insertion and securement with sterilized cotton threads.

After surgery, rats received a subcutaneous injection of saline solution (0.9% NaCl; 5–10 mL) for fluid replacement. During the 7-day recovery period, they were given daily subcutaneous Flunixin injections for at least 3 days and up to 5 days, depending on pain assessment using the Grimace scale (Mogil et al., 2020). Catheters were flushed daily with 0.2 mL of saline solution containing heparin (5000 IU/mL, Cristália) and gentamicin (15 mg/mL, Sigma-Aldrich), following a previously established protocol (Thrivikraman, Huot, & Plotsky, 2002).

### Cocaine self-administration and allocation to groups

The rats were approximately three months old at the start of the self-administration phase. We used self-administration chambers (Med Associates, St. Albans, VT, USA) equipped with two operant levers. One retractable on the right side (serving as the active lever in which responses resulted in cocaine delivery) and another non-retractable positioned on the left side, serving as the inactive lever. Pressing the active lever resulted in a 0.1 mL cocaine infusion over 2.3 seconds, accompanied by a 5-second presentation of discriminative cues (activation of a light above the lever and an auditory cue). The infusion delivery/active lever press ratio was fixed at 1:1, except during a 20-second timeout period that began immediately after cocaine delivery. During the timeout period, active lever responses were recorded but not reinforced. In addition, inactive lever presses were recorded but never reinforced. After connecting the catheter on the animal’s back to the self-administration system (an interface controlled by the MED_PC-IV Software, Med Associates®) each session began with the presentation of the active lever, activation of its corresponding light and auditory cue, and chamber illumination through an ambient light on the opposite wall, which remained on throughout cocaine availability. Additional visual cues, such as sticks on the cage walls, provided contextual stimuli.

Two cocaine self-administration protocols were implemented: restricted access (RA) and extended access (EA). RA sessions lasted 1 hour per day for 12 consecutive days. EA sessions followed the same structure but consisted of six 1-hour blocks separated by 5-minute intervals, resulting in a total session duration of 385 minutes. During these intervals, the active lever was retracted, and the ambient light was turned off. These conditions were based on previous studies of cue-induced incubation of cocaine craving (Grimm et al., 2003; Koya et al., 2009; Lu et al., 2004, 2007). Aiming to include only animals with relevant exposure to cocaine, during this phase we adopted the following inclusion criteria: self-administer a minimum of 5 infusions during the first hour on at least 3 out of the last 5 days of training. Forty-three rats (27 from RA; 16 from EA) were excluded from the next phases of the study due to these criteria. In addition to the number of infusions received and the number of presses on the active lever, cognitive performance was evaluated using the active/inactive lever press ratio as a discrimination index, while the infusion/active lever press ratio was used as an impulsivity indicator. A theoretical biological relevance threshold of 0.5 was applied, with values below this threshold indicating increased impulsivity-like behavior.

Following self-administration training, rats from both protocols (RA and EA) were subjected to either a 1-day or a 30-day forced abstinence period, during which they remained in their home cages without access to the self-administration chambers or cocaine. After their respective abstinence periods, rats were either kept in their home cages or re-exposed to the context in which self-administration had taken place. This experimental design resulted in eight groups as follows: RA-abstinence 1-home cage (R1-HC); RA-abstinence 1-context exposure (R1-CE); RA-abstinence 30-home cage (R30-HC); RA-abstinence 30-context exposure (R30-CE); EA-abstinence 1-home cage (E1-HC); EA-abstinence 1-context exposure (E1-CE); EA-abstinence 30-home cage (E30-HC); EA-abstinence 30-context exposure (E30-CE). Group assignment was conducted by first calculating (for each rat) the average number of active lever presses during the first hour of training over the last five days. These values were then arranged in ascending order and used to similarly distribute within-protocol training performance variability across groups.

### Re-exposure to drug environment and euthanasia

After 1 or 30 days of abstinence, animals were re-exposed to the self-administration environment, where they encountered the same contextual and discriminative cues previously associated with cocaine self-administration. The 30-minute test sessions did not include cocaine reinforcement; however, responses on the active lever triggered the presentation of discriminative cues. Ninety minutes after the start of re-exposure tests, animals were euthanized to capture the peak of Fos expression associated with neuronal activation induced by the drug context. Euthanasia was performed under anesthesia using ketamine (180 mg/kg), xylazine (30 mg/kg), and fentanyl (0.05 mg/kg), followed by transcardiac perfusion with 100 mL of 0.1M phosphate buffer (PB) and 200 mL of 4% paraformaldehyde in PB (PFA). The brains were removed, post-fixed in PFA for 30 minutes, and transferred to a 30% sucrose solution for cryoprotection over three days. They were then frozen in dry ice for 1 hour and stored at -80 °C until further processing for immunohistochemistry.

### Immunohistochemistry

The brains were sectioned into 30 µm slices and preserved in an antifreeze solution (20% glycerol + 30% ethylene glycol in 0.1M phosphate buffer). The slices were then placed in immunohistochemistry baskets and subjected to a washing step (five 5-minute incubation in PB). Antigen retrieval was performed by immersing the sections in a 0.01M sodium citrate solution at pH 8.00 for 20 minutes at 70 °C, followed by an additional 10 minutes of exposure at room temperature. After antigen retrieval, sections underwent a washing step and were incubated in a solution of 1% hydrogen peroxide diluted in PB for 20 minutes. After a washing step, sections were transferred to 2 ml plastic tubes for incubation with the primary antibody (mouse monoclonal Anti-c-Fos - ab208942, RRID: AB_2747772, Abcam, 1:2000; rabbit polyclonal Anti-Parvalbumin - Ab11427, RRID: AB_298032, Abcam, 1:3500; mouse monoclonal Anti-NeuN - MAB377, RRID: AB_2298772, Merck/Millipore, 1:2000). A 0.1M PB solution containing 1% Tween 20 (V/V) and 2% Molico™ powdered milk (M/M) was used for primary antibody incubation. This incubation occurred for 36 to 48 hours at 4-8 °C with slow agitation. Antibody concentrations were determined in preliminary assays. Following primary antibody incubation, sections underwent a washing step with subsequent incubation with secondary antibodies for 2 hours (goat anti-Mouse IgG, Alexa Fluor™ 488 - A32723, RRID: AB_2633275, 1:500; donkey anti-Mouse IgG, Alexa Fluor™ 568 - A10037, RRID: AB_11180865, 1:500; donkey anti-Rabbit IgG, Alexa Fluor™ 568 - A-21206, RRID: AB_2535792, 1:500. All provided by ThermoFisher Scientific) in PB with 1% Tween 20. All subsequent procedures were performed with reduced light to preserve fluorescence. After incubation, sections underwent three washes in PB, with DAPI (Sigma-Aldrich, D9542) added in the fourth wash at a concentration of 1:100,000. The sections were then washed twice with PB and transferred to slides. Following a brief drying period, slides were covered with coverslips using ProLong™ antifade mounting solution. Fluorescence images were acquired using a Zeiss Axio Imager D2™ fluorescence microscope equipped with a camera and controlled by Zen Blue (Carl Zeiss) software. The microscope light intensity was kept constant across all acquisitions, while the exposure time was automatically adjusted for each field of interest and each fluorescence channel. Each image covered an area of 0.145 mm², and no post-processing for signal enhancement was applied.

Images were exported in .tiff format and processed using QuPath. Before batch processing, representative images from different experimental groups were used to define an intensity threshold for signal detection in all three fluorescence channels (red, blue, and green). This threshold was subjectively determined by the experimenter and uniformly applied to all images. A QuPath macro was developed to automate cell detection based on the targeted labeling, quantifying the number of NeuN-positive cells per field and the subset that colocalized with Fos signal. The same analysis was performed for PV interneurons labeled with Fos.

### Bias-reducing measures and statistical analysis

In this study, we ran an exploratory experiment (no sample size calculation was performed in advance) in which each included rat was considered an experimental unit [pooled 71 rats from seven cohorts = C1 (n = 12); C2 (n = 10); C3 (n = 1); C4 (n = 8); C5 (n =16); C6 (n = 18); C7 (n = 6)]. No computed sequence generator was used to allocate animals to their respective interventions. However, selection bias was minimized by introducing similarly within-protocol baseline characteristics variability between groups, as described in the session “Cocaine self-administration and allocation to groups”. Performance bias was minimized by having a blinded assessor to digitize immunolabeled images. Detection bias was minimized using automated systems (an approach equivalent to blinding outcome assessors) to collect operant cocaine self-administration behavior data. A QuPath macro was developed to automate cell detection based on the targeted labeling, quantifying the number of NeuN-positive cells per field and the subset that colocalized with Fos signal. The same analysis was performed for PV interneurons labeled with Fos Attrition bias was minimized by adhering to all predefined exclusion criteria as follows: (i) identification of signs of pain or distress in the rats (no animals were excluded based on this criterion). (ii) cannula malpositioned or removed (11 from RA - 11.57%, and 15 from EA - 15.95%). (iii) poor performance on training (27 from RA - 32.14%, and 16 from EA - 20.25). (iv) specifically for discrimination index, not pressing the active lever at least once during re-exposure test (2 animals were excluded from this specific analysis during the contextual re-exposure test). (v) poor staining/image quality resulted in the exclusion of 3 animals from histological analysis. No outliers were excluded. Based on “ARRIVE guidelines 2.0: Essential 10 list” (Sert et al., 2020) available at https://arriveguidelines.org/arrive-guidelines and on “SYRCLE’s risk of bias tool for animal studies” (Hooijmans et al., 2014), all efforts have been made to ensure transparency and avoid reporting bias as recommended elsewhere (Loss et al., 2021).

Statistical analyses were performed following the recommendations described by Loss and colleagues (Loss et al., 2021). All data were analyzed in RStudio (Version 1.4.1717 – 2009– 2021 RStudio, PBC) R version 4.1.0 (2021–05-18) using the following strategy: (1) A hierarchical selection model approach (based on the Akaike Information Criterion corrected for small samples – AICc) evaluating all possible combinations of subsets of factors (through the dredge function from the MuMIn package, Bartoń, 2022) was applied to select the most parsimonious model (the model presenting the lowest AICc value), defined as the simplest model where the removal of any remaining predictors resulted in a significant loss of information. Selection model was applied on Generalized Linear Mixed Models (GLMM); (2) As an exploratory alternative, a GLMM was fitted separately for each group using “INF Training” (average of infusions of the first training hour during the last 5 days of training) as a predictor factor, and the resulting slope (beta coefficient and 95% confidence intervals) were plotted for the range values of “INF Training”. To control potential “cohort effect”, the experimental cohort (block) was included as a random factor in all GLMM. Based on (Schmettow, 2021), the family distribution and its canonical link function were chosen beforehand, considering the nature of the data. Specifically, continuous data presenting no upper boundaries but presenting lower boundaries = zero [mean number of active lever presses during training, discrimination index during training, percentage of Fos-positive cells (Fos/NeuN)] were analyzed using Gamma distribution with Log link [once Gamma distribution does not allow for non-positive values, when necessary (i.e., at least one y value = zero), a positive constant (epsilon = 1e-6) was added to the data set]; continuous data presenting both upper and lower boundaries (INF Training and % of Fos+ cells) were analyzed using Beta distribution with Logit link [once Beta distribution only allows values between zero and 1 (i.e., 0 < y < 1) data were analyzed as frequency; the constant epsilon was either added (when at least one y = 0) or subtracted (when at least one y = 1) to all data set]; discrete data (counts) presenting only lower boundaries = zero (number of active lever presses during test) were first analyzed using Poisson distribution with Log link followed by Negative Binomial distribution with Log link for the cases in which over-dispersion was detected during diagnosis procedure. All GLMM were performed using the glmmTMB package (Brooks et al., 2017). When necessary, the raw data were subjected to transformations before the analysis, as described in Supplementary Material.

During the training phase, only one fixed factor (protocol) and one random factor (cohort) were used. All selected models were submitted for diagnosis using the DHARMa package (Hartig et al., 2024). Except for overdispersion in the models using Poisson distribution (as described above), no action was made to avoid diagnosis problems (diagnosis outputs are described for each variable). A similar procedure was applied for the test phase except that three predictor variables were used (two categorical - factor 1: “protocol” and factor 2: “abstinence”; plus one continuous - factor 3: “INF Training”). For histological data, one additional categorical factor was added to the analysis (factor 4: “environment”). Thus, the full model (the most complex one) during histological analysis was [response variable ∼ protocol * abstinence * environment * INF Training + (1|cohort)]. For all experiments, estimated marginal means and 95 % confidence intervals (CI) were extracted from the selected model using the emmeans package (Lenth, 2022) and graph-plotted using GraphPad Prism 8. The presented statistical significance was based on the selected models. Bonferroni pairwise (post hoc) comparisons were performed for cases in which any interaction effect was detected, controlling for family-wise error. A two-tail 0.05 significance level (alpha) was set for all analyses.

## Results

### Training and maintenance of cocaine self-administration

In this phase, rats underwent twelve cocaine self-administration sessions. Performance was analyzed by averaging the first hour of the last five sessions. Both the RA and EA groups showed a progressive increase in cocaine infusions over time (i.e., escalation of cocaine consumption), accompanied by increased active lever presses and improved active/inactive lever discrimination. Selection model combined with GLMM revealed a protocol effect (EA > RA) for the average number of infusions (Figure 1a, Supplementary Table S1), active lever presses (Figure 1b, Supplementary Table S2), and active/inactive ratio (Figure 1c, Supplementary Table S3). These findings indicate that the escalation profile was more pronounced in the EA group, while it was discrete in the RA group. We further assessed the relationship between lever presses and infusions delivered as an indicator of impulsivity (here referred to as the impulsivity index). No effect was identified for this measure (i.e., the null model was selected) indicating no difference between groups for this variable as well as no relevant impulsivity-like behavior in both groups (Figure 1c, Supplementary Table S4). Of the 163 rats that completed self-administration training with adequate catheter patency, 120 met the inclusion criteria (i.e., receiving at least five infusions during the first hour in at least three of the last five training sessions). From the 43 rats that presented poor performance on training, 27 were from the RA group (i.e., 32% of rats in the RA group were excluded) while 16 were from EA group (i.e., 20% of rats in the EA group were excluded) indicating that the rats subjected to EA group had 17,5% more chances to be included than rats in the RA group (Fisher’s exact test, P= 0.1095; Relative Risk with 95% CI= 1.175; 0.9773 to 1.427). From the 120 rats that met the inclusion criteria, 71 of them were assigned to either undergoing a contextual Re-exposure session (RA1-CR: N=11, EA1-CR: N=10, RA30-CR: N=8, EA30-CR: N=9) or staying undisturbed in their home cages until the euthanasia (RA1-HC: N=8, EA1-HC: N=8, RA30-HC: N=7, EA30-HC: N=10). The remaining animals were allocated to other laboratory studies and were not included in the present experiments.

**Figure 1.**
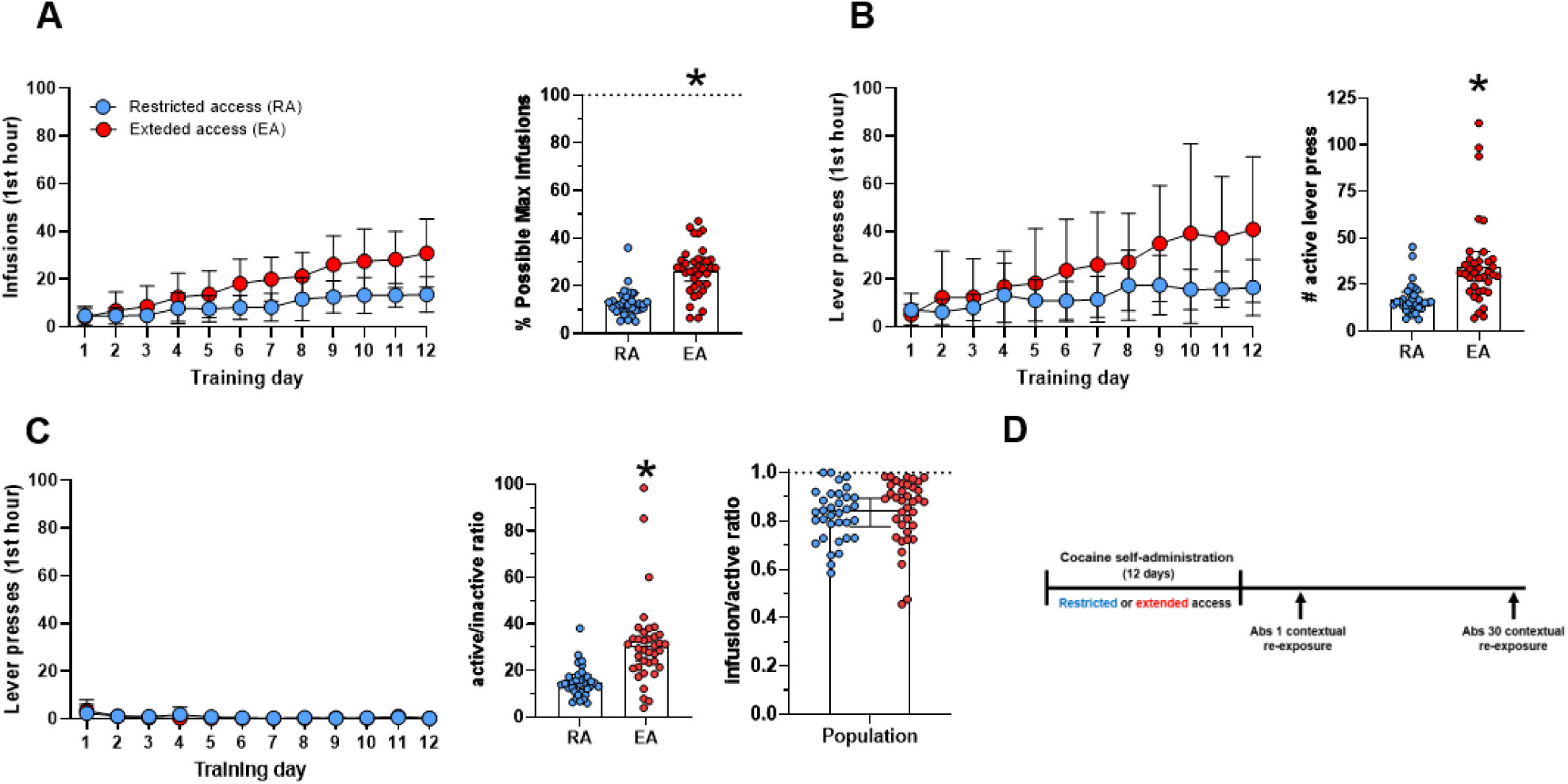
Cocaine intake escalates markedly during EA but remains discrete during RA. **A**, Number of cocaine infusions during the first training hour across 12 days of training (left) and the average number of infusions over the last 5 days of training expressed as a percentage of the maximum possible infusions during the first training hour (right); **B**, Number of active lever responses during the first training hour across 12 days of training (left) and the average number of active lever responses over the last 5 days of training (right); **C**, Number of inactive lever presses during the first training hour across 12 days of training (left), ratio of active to inactive lever presses (middle), and ratio of infusions to active lever presses (right) **D**, Experimental design. Following self-administration training rats were kept at their home-cage, and underwent forced abstinence for 1 or 30 days. They were then either euthanized directly from their home cage or after being re-exposed to the drug context in the presence of all contextual and discrete cues. No cocaine was delivered following active lever presses responses. Bar graph data were analyzed by GLMM and are presented as Estimated Marginal Means and their respective 95% CI. Individual values (observed units) are presented as dot-plots; * indicates p < 0.05 compared to RA; Line graphs’ data (presented as means ± SD) are representations of rats’ performance across time and were not subjected to statistical analysis. Only rats that met the training inclusion criteria are presented and no outliers were excluded (RA N = 34; EA N = 37).

### contextual re-exposure to cocaine self-administration environment

One or thirty days after the last training session (i.e., following an abstinence period of 1 or 30 days, respectively), rats from both the RA and EA groups were re-exposed to the context in which cocaine self-administration took place. However, responses on the active lever were no longer reinforced with cocaine infusions. Importantly, the average number of infusions during training (“INF Training”) can account for a portion of data variability that cannot be explained by the protocol factor alone, even that the protocols are nested within “INF Training” (i.e., “INF Training” for EA group > “INF Training” for RA group). Therefore, “INF Training” was also included as a predictor factor for all the GLMM conducted on post-training data.

For the number of active lever presses, a main abstinence effect was indicated by the selected model. GLMM analysis revealed that, independently of the protocol of training and “INF Training”, the number of active lever presses was higher in the 30-day compared to the 1-day abstinence period (Figure 2a). As we understand that the lack of interaction effect (either “abstinence * protocol” or “abstinence * INF Training”) might be due to low statistical power, we conducted an exploratory analysis to visualize the relationship between “INF Training” (as predictor factor) and number of active lever responses. This analysis consisted of fitting a GLMM separately for each group using only “INF Training” as a predictor factor. The resulting slopes indicated that for RA1, RA30 and EA30 (but not for EA1), the number of active lever presses during the re-exposure test increased as the “INF Training” increased (Supplementary Figure 2b). To further investigate this pattern, we performed three additional GLMMs. The selection of data to be included in these GLMMs were based either on the abstinence factor (i.e., 1-day groups were fit in one model and 30-day groups were fit in another model) or on the direction of the slopes from the group-specific GLMMs (i.e., classifying groups based on whether the relationship between “INF Training” and active lever pressing was positive or not). The “Model 1” was fitted with data from RA1 and EA1 altogether. Although non-statistically significant (P= 0.0665), this model might suggest that when the abstinence period is 1 day (i.e., for RA1 and EA1 data), the number of active lever presses during the contextual session decreases as the “INF Training” increases (Figure 2b); For the “Model 2”, fitted with data from RA30 and EA30 altogether - although non-statistically significant (P= 0.0561), it might suggest that when the abstinence period is 30 days (i.e., for RA30 and EA30 data), the number of active lever presses during the contextual session increases as the “INF Training” increases (Figure 2b); The “Model 3” - fitted using only the groups that presented positive slopes in the “group-specific GLMMs” approach (i.e., RA1, RA30 and EA30 were included in the model while, EA1 was not included) indicated that the number of active lever presses during the contextual session increased as the “INF Training” increased (P= 0.000175; Figure 2c). Taken together, it is possible to speculate that the abstinence period does not affect RA groups’ behavior in the context re-exposure test (i.e, RA1 and RA30 present similar profiles), but it affects EA groups behavior, supporting the notion that the EA protocol is more effective in inducing the incubation of cocaine craving. Additionally, we observed that while the EA30 group increased the number of active lever presses concerning its RA pair (RA30), the EA1 group presented a reduced number of active lever presses with its RA pair (RA1).

**Figure 2.**
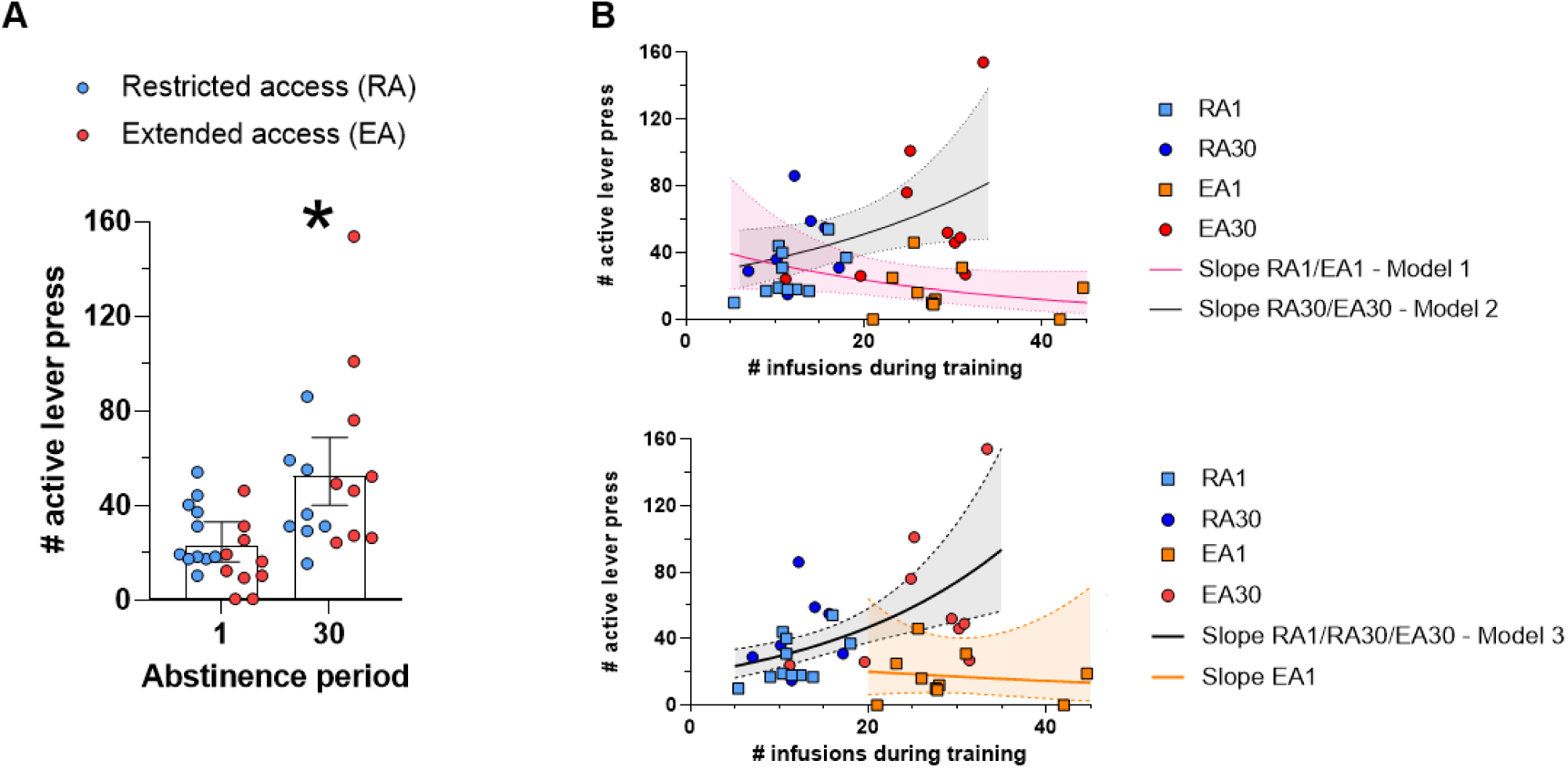
Cocaine intake during self-administration training modulates incubation of craving. **A**, Active lever responses measured during the 30-minute contextual re-exposure test following 1 or 30 days of abstinence; **B** Exploratory GLMM analysis of active lever responses as a function of cocaine intake during the last days of training (INF training). Three independent GLMM were fitted: (top panel) Model 1 grouped data from 1-day abstinence period (RA1 with EA1); Model 2 grouped data from 30-day abstinence period (RA30 with EA30); (bottom panel) based on the slopes resulting from the GLMM individually performed for each experimental group (see Supplementary Figure S5), Model 3 included groups presenting positive slopes (RA1, RA30, and EA30; EA1 was excluded). Bar graph data (in A) were analyzed by GLMM and are presented as Estimated Marginal Means and their respective 95% CI. Individual values (observed units) are presented as scatter-plots; * indicates p < 0.05 compared to 1 day of abstinence; In B, solid lines represent the Estimated Marginal Means while shaded area limited by dashed lines represent 95% CI. No outliers were excluded (RA N= 34; EA N= 37).

A similar approach was used for the active/inactive ratio analysis. No effect was identified in the model selection approach (i.e., the null model was selected) indicating no difference between groups for this variable (Figure 2a and Supplementary Table S6). As we understand that the lack of interaction effect (either “abstinence * Protocol” or “abstinence * INF Training”) might be a consequence of low statistical power, we performed an exploratory approach (as in the number of active lever presses) as follows: “Model 1” - indicated that when the abstinence period is 1 day (i.e., for RA1 and EA1 data), the active/inactive ratio reduces as the “INF Training” increases (P= 0.00497; RA1 mean = 21.3; 9.97-45.5 95% CI; EA1 mean = 6.1; 2.86-13.0 95% CI; Figure 3b); “Model 2” - indicated that “INF Training” does not influence the active/inactive ratio when the abstinence period is 30 days (i.e., for RA30 and EA30; P= 0.77; Population mean = 19.2; 12.3-30.0 95% CI; Figure 3b); “Model 3” - it was run using only RA1, RA30 and EA30 while EA1 was not included. Model 3 indicated that “INF Training” does not influence the active/inactive ratio for this data (P= 0.89; Population mean = 18.4; 12.4-27.1 95% CI; Figure 3b). Taken together, it is possible to speculate that the abstinence period does not affect RA groups active/inactive ratio in the context re-exposure test (i.e., RA1 and RA30 present similar profiles), but it affects EA groups active/inactive ratio. While the EA30 group presented a similar active/inactive ratio with its RA pair (RA30), the active/inactive ratio was reduced in the EA1 group with its RA pair (RA1).

**Figure 3.**
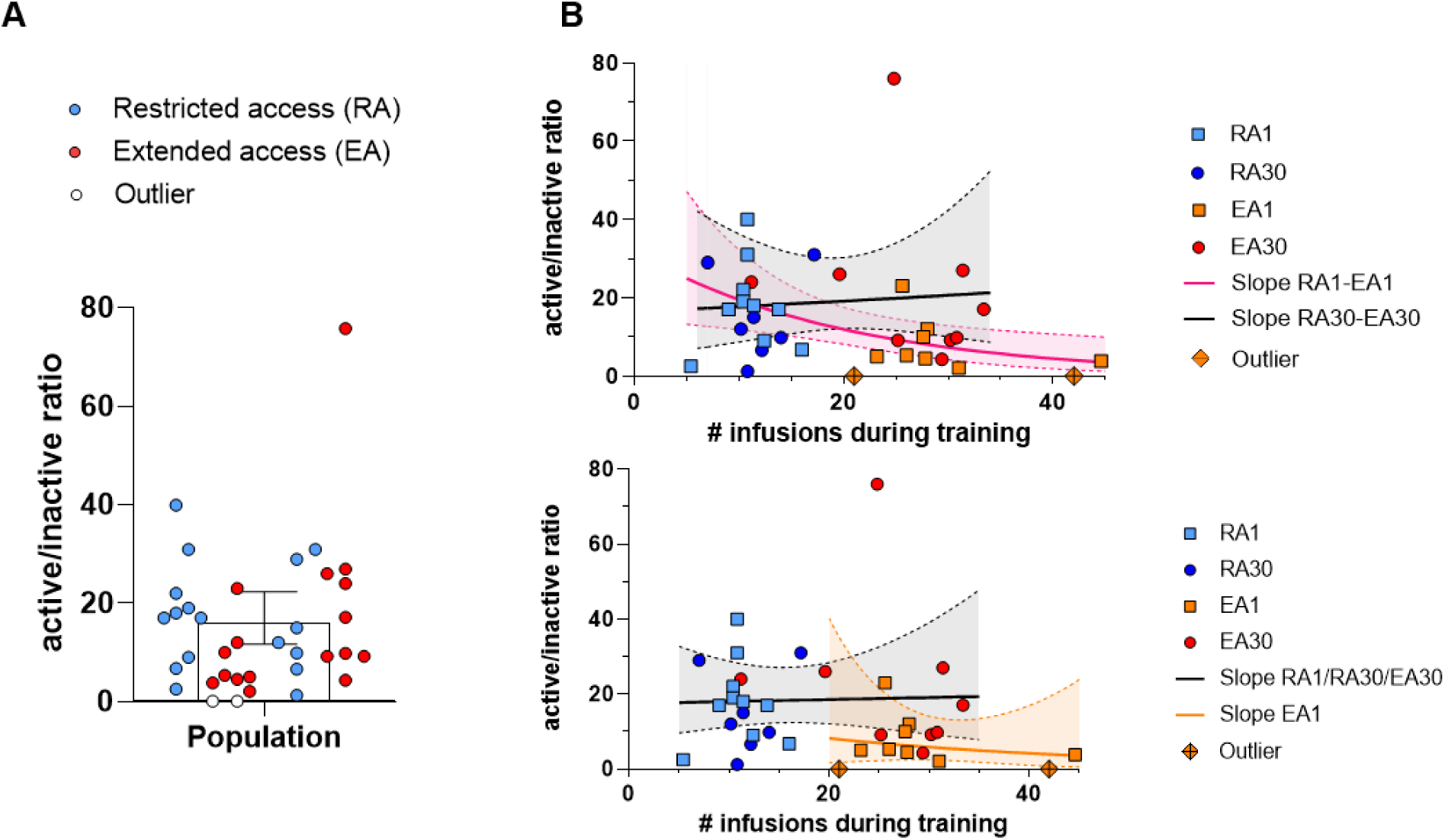
Cocaine intake during self-administration training differentially affects the active/inactive ratio following forced abstinence. **A**, Active/inactive ratio measured during the 30-minute contextual re-exposure test following 1 or 30 days of abstinence; **B**, Exploratory GLMM analysis of active/inactive ratio as a function of cocaine intake during the last days of training (INF training). The same approach used for active lever responses was applied for the active/inactive ratio analysis, as follows: three independent GLMM were fitted - (top panel) Model 1 grouped data from 1-day abstinence period (RA1 with EA1); Model 2 grouped data from 30-day abstinence period (RA30 with EA30); (bottom panel) Model 3 included RA1, RA30, and EA30 (EA1 was excluded). Bar graph data (in A) were analyzed by GLMM and are presented as Estimated Marginal Means and their respective 95% CI. Individual values (observed units) are presented as scatter-plots; In B, solid lines represent the Estimated Marginal Means while shaded area limited by dashed lines represent 95% CI. The two highlighted outliers presented in the graph were excluded from the analysis (RA1 N= 11; EA1 N= 8; RA30 N= 8; EA1 N= 9).

### Nucleus accumbens neuronal activation following re-exposure to the cocaine-associated context after RA or EA Self-Administration protocols and forced abstinence

Ninety minutes after the starting of the session (RA1-CR N=11; EA1-CR N=10; RA30-CR N=8; EA30-CR N=9), rats were euthanized and had their brains collected for histological evaluation. Home cage groups (RA1-HC N=8; EA1-HC N=8; RA30-HC N=7; EA30-HC N=10) were euthanized in a paired-matching way. For the the percentage of NeuN-positive cells that also expressed c-Fos (%Fos⁺/NeuN⁺) in NAcc Core, the selected model indicated an interaction effect between abstinence and environment, along with a main effect of “INF Training” (abstinence * Environment + INF Training) which was further confirmed by the GLMM analysis (Supplementary Table S7). Specifically, Bonferroni post hoc analysis indicated that %Fos+/NeuN+ was overall lower in home cage groups compared to the context re-exposed ones. While no between-groups difference was found for the home cage groups (i.e., RA1-HC = EA1-HC = RA30-HC = EA30-HC), for the context re-exposed ones there was a higher %Fos+/NeuN+ in the 30-day compared to the 1-day abstinence period groups (Figure 3 a, b, and c). In addition, we found that the %Fos+/NeuN+ increases as the “INF Training” increases (Figure 3 c - d). As previously mentioned, the Protocols are nested within the “INF Training” (i.e., “INF Training” for EA group > “INF Training” for RA group), meaning that the %Fos+/NeuN+ in NAcc Core was higher in the EA-CR compared to their respective RA-CR groups (i.e., EA1 > RA1; EA30 > RA30).

For the %Fos+/NeuN+ in NAcc Shell, the selected model indicated an interaction effect between abstinence and environment (abstinence * Environment) that was further confirmed in the GLMM analysis (Supplementary Table S8). Specifically, Bonferroni post hoc analysis indicated that %Fos+/NeuN+ was overall higher in the context re-exposed groups compared to the home cage ones. While no between-groups difference was found for the home cage groups (i.e., 1-day abstinence-HC = 30-day abstinence-HC), for the context re-exposed ones there was a higher %Fos+/NeuN+ in the 30-day compared to the 1-day abstinence period groups (Figure 3e - f). In contrast to the %Fos+/NeuN+ in NAcc Core analysis, neither the protocol nor the “INF Training” influenced the %Fos+/NeuN+ in NAcc Shell.

### Activation of parvalbumin positive interneurons from nucleus accumbens following re-exposure to the cocaine-associated context after RA or EA self-administration protocols and forced abstinence

We also quantified the percentage of parvalbumin interneurons expressing Fos under the same experimental conditions as the Fos+/NeuN+ labeling (RA1-CR N=4; EA1-CR N=5; RA30-CR N=5; EA30-CR N=4; RA1-HC N=5; EA1-HC N=6; RA30-HC N=6; EA30-HC N=7). However, not all samples in this experiment were obtained from the same rats used for Fos+/NeuN+ labeling. For the percentage of PV+ cells that also expressed c-Fos (%Fos+/PV+) in NAcc core, the selected model indicated a triple interaction effect between protocol, abstinence and environment (Protocol * abstinence * Environment) that was further confirmed in the GLMM analysis (Figure 4a - b and Supplementary Table S9). Regarding the home cage vs context re-exposure comparison, Bonferroni post hoc analysis indicated that the %Fos+/PV+ was higher in RA30-CR and EA1-CR in comparison to their home cage pairs (RA30-HC and EA1-HC, respectively) but not in RA1-CR or EA30-CR in comparison to their home cage pairs (RA1-HC and EA30-HC, respectively). In the comparison of the 30-day vs. 1-day abstinence periods for the context re-exposed groups, an increased percentage of %Fos+/PV+ was observed in the RA groups after 30 days of cocaine abstinence compared to the 1-day abstinence period (i.e., RA30-CR > RA1-CR). In contrast, the opposite effect was observed in the EA groups (i.e., EA30-CR < EA1-CR). Regarding the between-protocols comparison for the context re-exposed groups, while we observed an increased %Fos+/PV+ in the EA compared to the RA after a 1-day abstinence period, the opposite effect was found for the 30-day abstinence period (i.e., RA30-CR > EA30-CR). No between-group differences were found for the home cage groups either for the 30-day vs 1-day abstinence period comparison or for the between-protocols comparison, meaning that the %Fos+/PV+ was similar between RA1-HC, EA1-HC, RA30-HC and EA30-HC.

**Figure 4.**
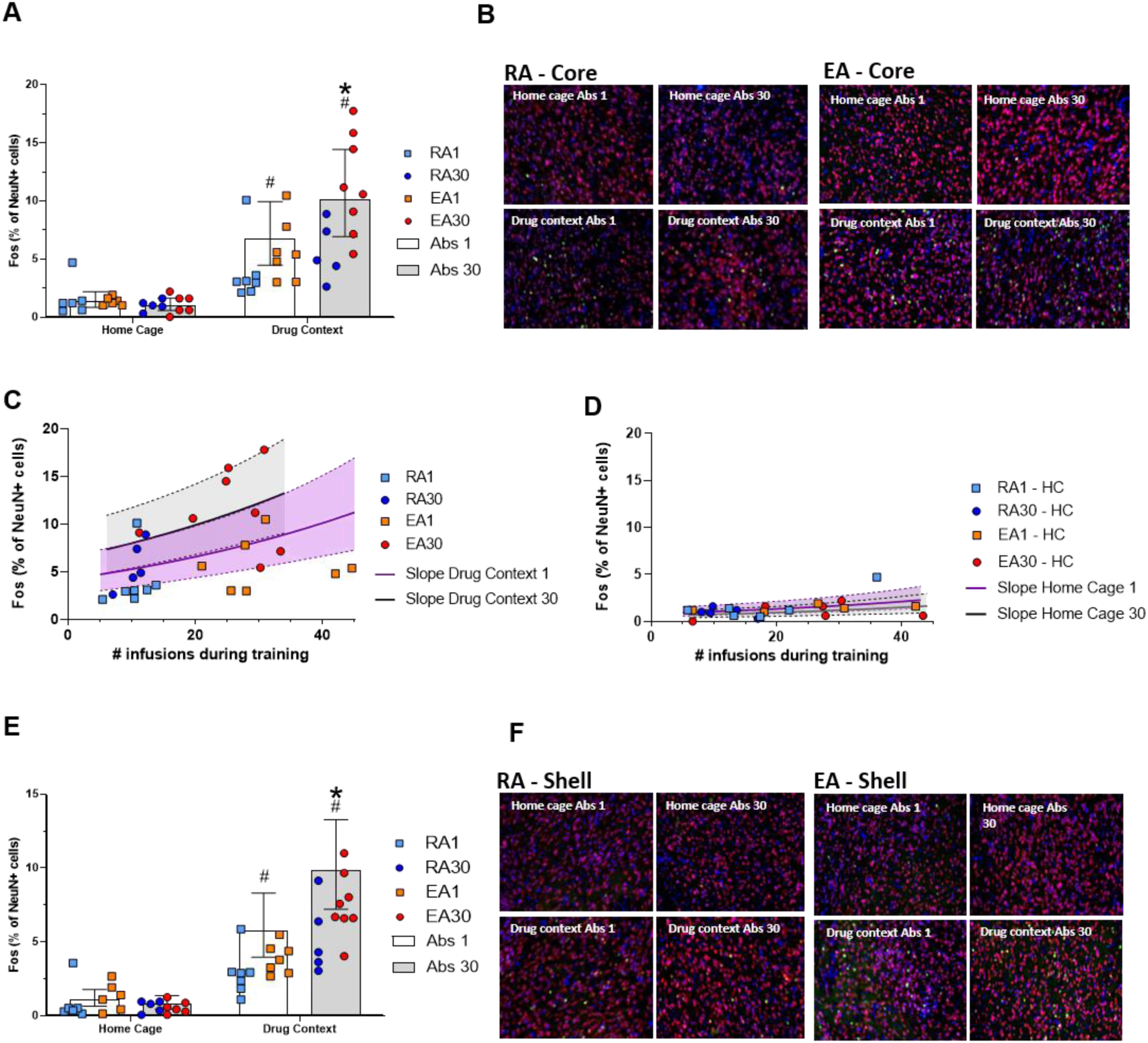
Neuronal activation in the nucleus accumbens core following drugcontext re-exposure increases with forced abstinence and cocaine intake. **A**, Percentage of NeuN-positive cells co-expressing Fos in the nucleus accumbens core after re-exposure to the drug-associated context or in rats not re-exposed to the drug context (Home Cage groups), following 1 or 30 days of abstinence; **B**, Representative image of NeuN (red) and Fos (green) staining in the NAcc core, with DAPI counterstain shown in blue; **C** and **D**, Neuronal activation in the NAcc core as a function of cocaine intake during the last days of training (INF training) after 1 or 30 days of abstinence for rats (C) that underwent drug-context re-exposure or (D) that were kept in their home cage without drug-context re-exposure; A-D together indicates that the neuronal activation in the NAcc core is higher in rats that underwent context re-exposure compared to rats that did not. Moreover, neuronal activation in the NAcc core is influenced by both cocaine intake during training and the duration of the abstinence period. Specifically, higher cocaine intake and longer abstinence periods (30 days) are associated with greater neuronal activation in the NAcc core. **E**, Percentage of NeuN-positive cells co-expressing Fos in the NAcc shell after re-exposure to the drug-associated context or in rats not re-exposed to the drug context (Home Cage group), following 1 or 30 days of abstinence; F, Representative image of NeuN (red) and Fos (green) staining in the NAcc shell, with DAPI counterstain shown in blue. All data were analyzed by GLMM and are presented as Estimated Marginal Means and their respective 95% CI. Individual values (observed units) are presented as scatter-plots; In C and D, solid lines represent the Estimated Marginal Means while shaded area limited by dashed lines represent 95% CI. * indicates p < 0.05 compared to 1 day of abstinence in the same environment; # indicates p < 0.05 compared to home cage environment in the same day of abstinence; No outliers were excluded from the analysis (RA1-HC N=8; EA1-HC N=8; RA30-HC N=7; EA30-HC N=10 RA1-CR N=11; EA1-CR N=10; RA30-CR N=8; EA30-CR N=9).

**Figure 5.**
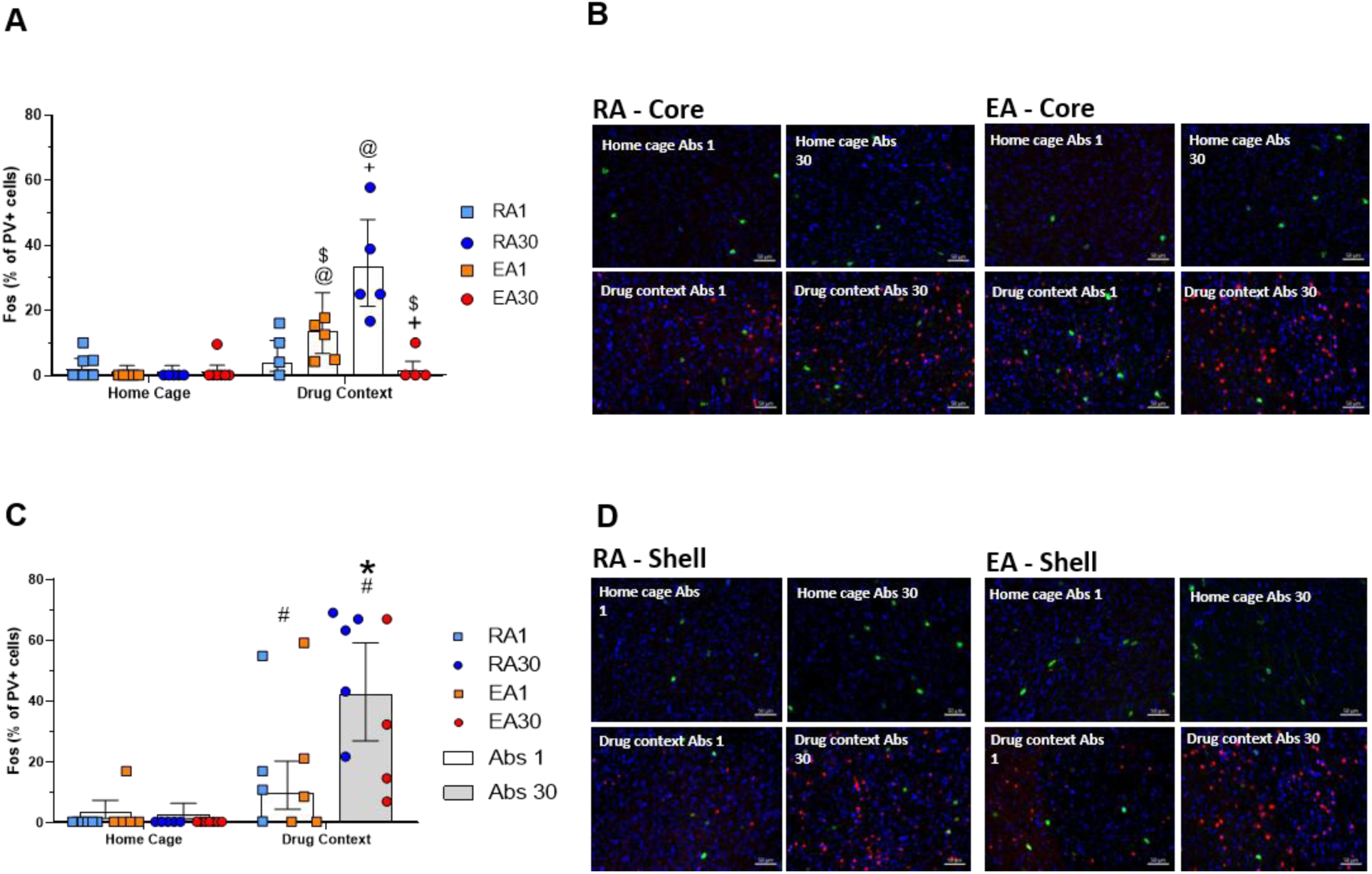
Decreased PV activation in the Nacc core following forced abstinence only in EA rats. **A**, Percentage of PV-positive cells co-expressing Fos in the nucleus accumbens core after re-exposure to the drug-associated context following 1 or 30 days of abstinence, or in rats not re-exposed to the drug context (Home Cage groups); **B**, Representative image of PV (green) and Fos (red) staining in the NAcc core, with DAPI counterstain shown in blue; **C**, Percentage of PV-positive cells co-expressing Fos in the nucleus accumbens shell after re-exposure to the drug-associated context, or in rats not re-exposed to the drug context (Home Cage groups) following 1 or 30 days of abstinence; **D**, Representative image of PV (green) and Fos (red) staining in the NAcc shell, with DAPI counterstain shown in blue. All data were analyzed by GLMM and are presented as Estimated Marginal Means and their respective 95% CI. Individual values (observed units) are presented as scatter-plots; @ indicates p < 0.05 compared to home cage environment in the same protocol and day of abstinence; $ indicates p < 0.05 compared to 1 day of abstinence in the same environment and protocol; + indicates p < 0.05 compared to RA in the same environment and abstinence period; * indicates p < 0.05 compared to 1 day of abstinence in the same environment; # indicates p < 0.05 compared to home cage environment in the same day of abstinence; No outliers were excluded from the analysis (RA1-CR N=4; EA1-CR N=5; RA30-CR N=5; EA30-CR N=4; RA1-HC N=5; EA1-HC N=6; RA30-HC N=6; EA30-HC N=7).

For the percentage of PV+ cells that are also Fos+ (%Fos+/PV+) in NAcc Shell, the selected model indicated an interaction effect between abstinence and Environment (abstinence * Environment) that was further confirmed in the GLMM analysis (Supplementary Table S10). Specifically, Bonferroni post hoc analysis indicated that %Fos+/PV+ was overall lower in home cage groups compared to the context re-exposed ones. While no between-groups difference was found for the home cage groups (i.e., 1-day abstinence-HC = 30-day abstinence-HC), for the context re-exposed ones there was a higher %Fos+/PV+ in the 30-day compared to the 1-day abstinence period groups (Figure 4c - d). In contrast to the %Fos+/PV+ in NAcc Core analysis, neither the Protocol nor the “INF Training” influenced the %Fos+/PV+ in NAcc Shell.

## Discussion

Our findings suggest that the regimen of cocaine self-administration, and consequently the cocaine intake, influences the self-administration profile, incubation of cocaine craving, and NAcc neuronal activation following drug-context re-exposure. Specifically, incubation of cocaine craving was associated with decreased activation of PV interneurons.

Both restricted and extended access protocols effectively induced self-administration learning, demonstrating that a one-hour daily exposure to cocaine is sufficient to establish associative learning. Notably, restricted access rats had a daily average cocaine consumption of 6.5 mg/kg of body weight during the last five days of training. Their average body weight of 320 g corresponds to an individual total intake of approximately 2 mg per session. This dose is relatively low compared to other studies using restricted access, yet it was sufficient to induce self-administration (Kawa et al., 2016). Furthermore, escalation in cocaine seeking and intake was observed in both protocols; however, this escalation was more pronounced in the extended access group, while it remained relatively moderate in the restricted access group.

We also observed that extended-access rats exhibited a discrimination index ten times higher than that of restricted-access rats. However, this difference was primarily driven by a progressive increase in active lever responses in the extended access group rather than by differences in discrimination or learning per se. Both groups showed reduced inactive lever presses during the first training days and maintained these values at consistently low levels thereafter, indicating comparable learning.

Regarding the impulsivity index, no significant differences were observed between groups. The impulsivity index was calculated as the ratio of infusions to the number of active lever presses, where values closer to 1 indicate a similar rate of active lever pressing and infusions received. A lower discrimination index can be interpreted as an indicator of craving, meaning that rats persist in seeking the drug despite its temporary unavailability. Alternatively, it may reflect impaired learning, as rats continue to use a suboptimal strategy, pressing the lever multiple times for a single infusion instead of pressing only once.

In our results, restricted and extended access protocols yielded an impulsivity index of approximately 0.9. When considered alongside the discrimination index, these data suggest that both groups developed a satisfactory learning curve, ultimately establishing an optimized drug-seeking behavior. However, it is important to highlight again that escalation was consistently higher in the extended access group.

Our findings, in line with existing literature, reinforce the evidence that different self-administration protocols and drug availability can shape consumption patterns and behavioral phenotypes (Ahmed & Koob, 1998; Guillem et al., 2014). Consistent with previous studies by Ahmed and colleagues, our results highlight the value of comparing restricted and extended access protocols as a tool for investigating the transition from controlled to compulsive drug use (Ahmed, 2010; Ahmed et al., 2002; Ahmed & Koob, 1998, 1999, 2004).

In humans, the transition from occasional use to substance use disorder is marked by a gradual escalation in drug consumption. Observing this escalation in animal models is a key predictor of a model’s ability to replicate the shift from recreational to compulsive drug use (Ahmed et al., 2000; Ahmed & Koob, 1998; Deroche-Gamonet et al., 2004; Deroche-Gamonet & Piazza, 2014; Lüscher et al., 2020). Escalation is often viewed as a consequence of behavioral changes linked to addiction, such as increased motivation to obtain the drug and a diminished capacity to limit its intake. However, this phenomenon should be interpreted with caution, as not all animals that escalate consumption will continue drug use despite punishment. According to Piazza and colleagues (2004), this persistence in the face of negative consequences is a critical behavior for identifying predictors of substance use disorder (Deroche-Gamonet et al., 2004; Lüscher et al., 2020; Pelloux et al., 2007).

Importantly, escalation is not dependent on sensitization or tolerance processes but is linked to alterations in motivational and hedonic mechanisms underlying compulsive behavior. While it is not the sole predictor of a model’s validity, it remains one of the most relevant (Ahmed, 2011; Ahmed et al., 2002; Ahmed & Cador, 2006; Ahmed & Koob, 1998; Paterson & Markou, 2003; Wee et al., 2007).

Beyond escalation, other behavioral evidence supports the extended access protocol as a more robust model for studying substance use disorders. For instance, rats under extended access persist longer in drug-seeking behavior even when the behavior is no longer reinforced. In progressive ratio assessments, extended access rats are more motivated to obtain the drug than restricted access rats. Furthermore, extended access rats are more likely to disregard potential danger and take significant risks to get the substance. They also tend to be more sensitive to relapse triggered by priming or stress compared to those under restricted access (Ahmed et al., 2000, 2002; Ahmed & Koob, 1998, 2004; Guillem et al., 2014; Vanderschuren & Everitt, 2004; Wolf, 2016).

Rats subjected to the extended access protocol consume significantly higher amounts of cocaine compared to those with restricted access. This has shifted discussions regarding the importance of the quantity of cocaine consumed in driving neurobiological changes associated with escalation and increased motivation for drug seeking and taking(Ahmed et al., 2002; Ahmed & Koob, 1998, 1999; Paterson & Markou, 2003). In this context, Kawa et al. (2016) proposed that an intermittent cocaine self-administration regimen, resulting in much lower consumption compared to extended access, could still induce compulsive behavior in rats, with marked escalation. In this study, animals underwent daily 4-hour cocaine self-administration sessions, with 5-minute self-administration periods alternated by 25-minute drug-free intervals. The study suggested that fluctuations in plasma cocaine levels and, consequently, dopamine levels in the synaptic cleft play a more critical role in the escalation of consumption and the development of compulsive behavior than the total amount of cocaine consumed.

In our study, the total daily cocaine consumption during late post-escalation sessions in the restricted access protocol was consistently lower than that reported by Kawa, Bentzley, and Robinson (2016), with rats consuming approximately 2 mg in our study compared to around 10 mg in theirs. This finding contributes to the ongoing discussion by suggesting that, in addition to fluctuations in plasma cocaine levels, a minimal level of cocaine consumption may also be necessary to induce a compulsive-like phenotype. Furthermore, it is important to note that in the extended access protocol, despite prolonged free access to the drug, consumption is not always stable, and fluctuations in plasma cocaine levels can also be present in this protocol.

We further investigated how different cocaine self-administration regimens influence the incubation of cocaine craving. As previously described, we measured the number of active lever presses during context re-exposure tests (Pickens et al., 2011). For both cocaine access regimens, the 30-day abstinence period increased drug craving. However, our data suggest that cocaine craving during contextual re-exposure tests intensifies with increased cocaine intake during self-administration training. Consequently, since rats in the extended access protocol consume more cocaine, they are more likely to exhibit the incubation of cocaine craving.

Interestingly, the analysis of the interaction between the number of infusions during training and the number of active lever presses during drug context re-exposure reveals that, after 30 days of forced abstinence, restricted access rats maintain a drug-seeking pattern similar to that observed before abstinence (Figure 2b). In contrast, the effects of forced abstinence on extended-access rats are much more pronounced. It is worth noting that extended access rats exhibited significantly lower cravings during the test conducted after one day of abstinence. This could be due to their high cocaine intake during the final training session, combined with the relatively short interval between the last session and the test (∼18 hours), which may have led to satiety and a temporary suppression of craving. These findings underscore the importance of cocaine intake in shaping the compulsive phenotype and highlight the interaction between minimal drug intake and abstinence in driving the incubation of cocaine craving, a key marker of compulsive drug-taking behavior (Deroche-Gamonet et al., 2004; Deroche-Gamonet & Piazza, 2014; Paterson & Markou, 2003).

Our results revealed that neuronal activation in the NAcc core increased after thirty days of forced abstinence in both restricted and extended access rats. Interestingly, this heightened responsiveness was context-specific, occurring only during drug-related re-exposure and not in a generalized manner (e.g., when the animals were in their home cages). However, regardless of whether abstinence lasted one or thirty days, NAcc activation remained consistently higher in extended access rats compared to restricted access rats. Notably, the level of NAcc activation in both groups was directly influenced by the amount of cocaine consumed during training, with this pattern mirroring the drug-seeking behavior observed during the contextual re-exposure tests. In parallel, restricted access rats showed increased PV (parvalbumin) activation after thirty days of abstinence, while extended access rats exhibited a marked reduction in PV activation after the same period.

Regarding the NAcc shell, thirty days of forced abstinence also induced higher neuronal activation following drug-related contextual re-exposure in both restricted and extended access rats. However, no differences in activation were observed between the protocols, and the amount of cocaine consumed during training did not influence this activation. For both restricted and extended access, PV activation was also higher following thirty days of abstinence.

Based on these findings, we speculate that the transition from occasional to compulsive drug-taking may be driven by molecular alterations in the NAcc core that enhance its responsiveness to drug-related cues. Additionally, the heightened activation of the NAcc core following forced abstinence could be linked to impaired inhibitory control of NAcc core MSNs by PV interneurons (Berke, 2011; Hu et al., 2014; Koós & Tepper, 1999). This idea aligns with experimental evidence indicating that increased neuronal reactivity in the NAcc, driven by enhanced glutamatergic transmission in response to cocaine cues, is ultimately necessary for the expression of incubated craving (Conrad et al., 2008; Dong et al., 2017). Furthermore, the NAcc core is crucial for instigating approach behavior towards reward-predictive cues. Exposure to a previously drug-paired environment can elicit a surge in glutamate release within the NAcc Core, triggering reinstatement of drug-seeking behaviors even after prolonged abstinence (Scofield et al., 2016; Thomas et al., 2009).

The overall increase in NAcc activation after abstinence in both protocols suggests that this activation may serve, to some extent, as a marker of associative learning between context and drug-seeking behavior. However, this does not necessarily indicate compulsivity, as NAcc core activity typically rises in response to reward-predictive cues and correlates with the vigor of approach behavior. Notably, the even greater activation observed in the NAcc core—both after one and thirty days of abstinence—along with the reduction in PV activation, may underlie the transition from occasional to compulsive drug-taking and the incubation of cocaine craving.

While there is substantial evidence linking PV interneurons to psychostimulant-induced behaviors, specific studies directly connecting these interneurons to the incubation of cocaine craving are limited. However, given their role in modulating MSN activity and influencing circuit adaptations, it is plausible that PV interneurons contribute to the neural mechanisms underlying incubation of cocaine craving. In this context, our work provides new evidence supporting the involvement of PV interneurons in the incubation of cocaine craving (Alexander 2022). However, further results are needed to establish a causal relationship between the disruption of PV interneuron inhibitory control and the incubation of cocaine craving. Moreover, it is important to clarify which populations of MSNs in the NAcc core, either D1 or D2, are more activated following forced abstinence in extended access rats.

Substance use does not equate to substance use disorder. While consumption is a key factor in the disorder’s development, these two patterns of use exhibit distinct behavioral and neurobiological characteristics (Lüscher et al., 2020). Therefore, animal models should account for this distinction by incorporating behavioral validations that extend beyond mere substance administration (Ahmed, 2010; Ahmed & Koob, 1998).

In our study, we provide new evidence highlighting the value of comparing different protocols to better understand the mechanisms underlying the transition from occasional to compulsive drug taking, the incubation of cocaine craving, and the circuit adaptations involved in this process. By refining our understanding of these mechanisms, our findings offer valuable insights into the ongoing discussion about animal models and their translational relevance in research on human substance use disorders.

## Acknowledgements

This work was supported by funds from São Paulo Research Foundation FAPESP (2018/15505-5, 2018/14153-7 and 2021/07134-9) and Coordenação de Aperfeiçoamento de Pessoal de Nível Superior (CAPES).

