## Supplementary tables for "Differential Effects of Cocaine Self-Administration Regimens on Incubation of Cocaine Craving and Nucleus Accumbens Neuronal Ensembles Activated by Cocaine-Associated Context"

### SUPPLEMENTARY MATERIAL

**Supplementary Table S1:** Summary of the selected model for analysis of infusions during the first hour of training.

| | | | | | |
|---|---|---|---|---|---|
| **Method** | Generalized Linear Mixed Model (GLMM) fit by maximum likelihood | | | | |
| **Family distribution with link function** | Beta ( logit ) | | | | |
| **Formula** | response_var ~ Protocol + (1 \| Cohort) | | | | |
| **Random effects** | | **Variance** | **SD** | | |
| | Cohort (intercept) | 0.03782 | 0.1945 | | |
| **# Observations** | 71 | | | | |
| **# Cohort** | 7 | | | | |
| **Fixed effects** | | **β** | **SE** | **t value** | **Pr(>\|z\|)** |
| | Intercept | -1.8505 | 0.1263 | -14.652 | <2e-16*** |
| | Protocol2Extended | 0.7896 | 0.1280 | 5.6.167 | 6.96e-10** |
| **Diagnosis** | No diagnosis problem detected. | | | | |
| **Other information** | | | | | |

Intercept = Population. SD = Standard Deviation; β = estimated coefficients for the fixed effect terms; SE = Standard Error. GLMM: Generalized Linear Mixed Model; Significance codes: *** 0.001; ** 0.01; * 0.05.

**Supplementary Table S2:** Summary of the selected model for analysis of active lever responses during the first hour of training.

| | | | | | |
|---|---|---|---|---|---|
| **Method** | Generalized Linear Mixed Model (GLMM) fit by maximum likelihood | | | | |
| **Family distribution with link function** | Gamma ( log ) | | | | |
| **Formula** | response_var ~ Protocol + (1 \| Cohort) | | | | |
| **Random effects** | | **Variance** | **SD** | | |
| | Cohort (intercept) | 0.03051 | 0.1747 | | |
| **# Observations** | 71 | | | | |
| **# Cohort** | 7 | | | | |
| **Fixed effects** | | **β** | **SE** | **t value** | **Pr(>\|z\|)** |
| | Intercept | 2.8171 | 0.1150 | 24.25 | <2e-16*** |
| | Protocol2Extended | 0.7146 | 0.1289 | 5.542 | 2.098e-08*** |
| **Diagnosis** | No diagnosis problem detected. | | | | |
| **Other information** | | | | | |

Intercept = Population. SD = Standard Deviation; β = estimated coefficients for the fixed effect terms; SE = Standard Error. GLMM: Generalized Linear Mixed Model; Significance codes: *** 0.001; ** 0.01; * 0.05.

**Supplementary Table S3:** Summary of the selected model for analysis of active/inactive ratio during the first hour of training.

| | | | | | |
|---|---|---|---|---|---|
| **Method** | Generalized Linear Mixed Model (GLMM) fit by maximum likelihood | | | | |
| **Family distribution with link function** | Gamma ( log ) | | | | |
| **Formula** | response_var ~ Protocol + (1 \| Cohort) | | | | |
| **Random effects** | | **Variance** | **SD** | | |
| | Cohort (intercept) | 0.03546 | 0.1883 | | |
| **# Observations** | 71 | | | | |
| **# Cohort** | 7 | | | | |
| **Fixed effects** | | **β** | **SE** | **t value** | **Pr(>\|z\|)** |
| | Intercept | 2.6862 | 0.1152 | 23.308 | <2e-16*** |
| | Protocol2Extended | 0.7628 | 0.1210 | 6.006 | 1.91e-09*** |
| **Diagnosis** | No diagnosis problem detected. | | | | |
| **Other information** | | | | | |

Intercept = Population. SD = Standard Deviation; β = estimated coefficients for the fixed effect terms; SE = Standard Error. GLMM: Generalized Linear Mixed Model; Significance codes: *** 0.001; ** 0.01; * 0.05.

**Supplementary Table S4:** Summary of the selected model for analysis of infusion/active ratio during the first hour of training.

| | | | | | |
|---|---|---|---|---|---|
| **Method** | Generalized Linear Mixed Model (GLMM) fit by maximum likelihood | | | | |
| **Family distribution with link function** | Beta ( logit ) | | | | |
| **Formula** | response_var ~ Protocol + (1 \| Cohort) | | | | |
| **Random effects** | | **Variance** | **SD** | | |
| | Cohort (intercept) | 0.2296 | 0.4792 | | |
| **# Observations** | 71 | | | | |
| **# Cohort** | 7 | | | | |
| **Fixed effects** | | **β** | **SE** | **t value** | **Pr(>\|z\|)** |
| | Intercept | 1.6962 | 0.2246 | 7.554 | 4.23e-14*** |
| **Diagnosis** | No diagnosis problem detected. | | | | |
| **Other information** | for BETA DISTRIBUTION the variable was transformed to 0 < y < 1, so the data fits the Beta distribution.<br>For the cases in which "variable1" has values = to the upper limit with all values > lower limit: a negative constant (epsilon) was added to data.<br>For the cases in which "variable1" has values = to the lower limit with all values < upper limit: a positive constant (epsilon) was added to data.<br>For the cases in which "variable1" has values = to the upper limit and also to the lower limit: epsilon (either negative or positive) was added only to values = to the limit data.<br>epsilon <- 1e-6 | | | | |

Intercept = Population. SD = Standard Deviation; β = estimated coefficients for the fixed effect terms; SE = Standard Error. GLMM: Generalized Linear Mixed Model; Significance codes: *** 0.001; ** 0.01; * 0.05.

**Supplementary Table S5:** Summary of the selected model for analysis of the number of active lever responses during the contextual re-exposure test.

| | | | | | |
|---|---|---|---|---|---|
| **Method** | Generalized Linear Mixed Model (GLMM) fit by maximum likelihood | | | | |
| **Family distribution with link function** | nbinom1 ( log ) | | | | |
| **Formula** | response_var ~ Abstinence + (1 \| Cohort) | | | | |
| **Random effects** | | **Variance** | **SD** | | |
| | Cohort (intercept) | 1.021e-09 | 3.19e-05 | | |
| **# Observations** | 38 | | | | |
| **# Cohort** | 7 | | | | |
| **Fixed effects** | | **β** | **SE** | **t value** | **Pr(>\|z\|)** |
| | Intercept | 3.1446 | 0.1753 | 17.935 | <2e-16*** |
| | Abs30 | 0.8157 | 0.2096 | 3.891 | 9.99e-05*** |
| **Diagnosis** | No diagnosis problem detected. | | | | |
| **Other information** | Intervals were back-transformed from the log scale. | | | | |

Intercept = Population. SD = Standard Deviation; β = estimated coefficients for the fixed effect terms; SE = Standard Error. GLMM: Generalized Linear Mixed Model; Significance codes: *** 0.001; ** 0.01; * 0.05.

**Supplementary Table S6:** Summary of the selected model for analysis of the active/inactive ratio during the contextual re-exposure test.

| | | | | | |
|---|---|---|---|---|---|
| **Method** | Generalized Linear Mixed Model (GLMM) fit by maximum likelihood | | | | |
| **Family distribution with link function** | Gamma ( log ) | | | | |
| **Formula** | response_var ~ 1 + (1 \| Cohort) | | | | |
| **Random effects** | | **Variance** | **SD** | | |
| | Cohort (intercept) | 0.01885 | 0.1373 | | |
| **# Observations** | 34 | | | | |
| **# Cohort** | 6 | | | | |
| **Fixed effects** | | **β** | **SE** | **t value** | **Pr(>\|z\|)** |
| | Intercept | 2.7802 | 0.1592 | 17.46 | <2e-16*** |
| **Diagnosis** | No diagnosis problem detected. | | | | |
| **Other information** | | | | | |

Intercept = Population. SD = Standard Deviation; β = estimated coefficients for the fixed effect terms; SE = Standard Error. GLMM: Generalized Linear Mixed Model; Significance codes: *** 0.001; ** 0.01; * 0.05.

**Supplementary Table S7:** Summary of the selected model for analysis of the percentage of NeuN cells co-expressing Fos in nucleus accumbens core

| | | | | | |
|---|---|---|---|---|---|
| **Method** | Generalized Linear Mixed Model (GLMM) fit by maximum likelihood | | | | |
| **Family distribution with link function** | Beta ( logit ) | | | | |
| **Formula** | response_var ~ Abstinence * Environment + Infusion_1h_Training + (1 \| Batch) | | | | |
| **Random effects** | | **Variance** | **SD** | | |
| | Cohort (intercept) | 0.2488 | 0.4988 | | |
| **# Observations** | 50 | | | | |
| **# Cohort** | 7 | | | | |
| **Fixed effects** | | **β** | **SE** | **t value** | **Pr(>\|z\|)** |
| | Intercept | -3.118548 | 0.24125 | -12.927 | <2e-16*** |
| | Abs30 | 0.444597 | 0.10781 | 4.124 | 3.73e-05 *** |
| | Home Cage | -1.648858 | 0.17554 | -9.393 | <2e-16*** |
| | INF Training | 0.023386 | 0.00481 | 4.856 | 1.20e-06 *** |
| | Abs30: Home Cage | -0.792707 | 0.25399 | 3.121 | 0.0018 ** |
| **Diagnosis** | No diagnosis problem detected. | | | | |
| **Other information** | | | | | |

Intercept = Population. SD = Standard Deviation; β = estimated coefficients for the fixed effect terms; SE = Standard Error. GLMM: Generalized Linear Mixed Model; Significance codes: *** 0.001; ** 0.01; * 0.05.

**Supplementary Table S8:** Summary of the selected model for analysis of the percentage of NeuN cells co-expressing Fos in nucleus accumbens shell.

| | | | | | |
|---|---|---|---|---|---|
| **Method** | Generalized Linear Mixed Model (GLMM) fit by maximum likelihood | | | | |
| **Family distribution with link function** | Beta ( logit ) | | | | |
| **Formula** | response_var ~ Abstinence * Environment + (1 \| Batch) | | | | |
| **Random effects** | | **Variance** | **SD** | | |
| | Cohort (intercept) | 0.1094 | 0.3307 | | |
| **# Observations** | 50 | | | | |
| **# Cohort** | 7 | | | | |
| **Fixed effects** | | **β** | **SE** | **t value** | **Pr(>\|z\|)** |
| | Intercept | -2.7947 | 0.1957 | 14.277 | <2e-16*** |
| | Abs30 | 0.5794 | 0.1447 | 4.005 | 6.21e-05 *** |
| | Home Cage | -1.7293 | 0.2490 | -6.945 | <2e-16*** |
| | Abs30: Home Cage | -0.9028 | 0.3473 | -2.600 | 0.00933 ** |
| **Diagnosis** | No diagnosis problem detected. | | | | |
| **Other information** | | | | | |

Intercept = Population. SD = Standard Deviation; β = estimated coefficients for the fixed effect terms; SE = Standard Error. GLMM: Generalized Linear Mixed Model; Significance codes: *** 0.001; ** 0.01; * 0.05.

**Supplementary Table S9:** Summary of the selected model for analysis of the percentage of PV cells co-expressing Fos in nucleus accumbens core.

| | | | | | |
|---|---|---|---|---|---|
| **Method** | Generalized Linear Mixed Model (GLMM) fit by maximum likelihood | | | | |
| **Family distribution with link function** | Beta ( logit ) | | | | |
| **Formula** | response_var ~ Protocol * Abstinence * Environment + (1 \| Batch) | | | | |
| **Random effects** | | **Variance** | **SD** | | |
| | Cohort (intercept) | 3.503e-09 | 5.919e-05 | | |
| **# Observations** | 41 | | | | |
| **# Cohort** | 4 | | | | |
| **Fixed effects** | | **β** | **SE** | **t value** | **Pr(>\|z\|)** |
| | Intercept | -3.2163 | 0.1957 | 14.277 | <2e-16*** |
| | Protocol2Extended | 1.3694 | 0.6250 | 2.191 | 0.028439 * |
| | Abs30 | 2.5234 | 0.6107 | 4.132 | 3.60e-05 *** |
| | Home Cage | -0.6857 | 0.6381 | -1.075 | 0.282591 |
| | Protocol2Extended: Abs30 | -4.9424 | 0.9370 | -5.275 | 1.33e-07 *** |
| | Protocol2Extended: Home Cage | -2.0284 | 0.8754 | -2.317 | 0.020491 * |
| | Abs30:Home Cage | -3.1824 | 0.8676 | -3.668 | 0.000244 *** |
| | Protocol2Extended: Abs30:Home Cage | 5.7588 | 1.2717 | 4.528 | 5.94e-06 *** |
| **Diagnosis** | No diagnosis problem detected. | | | | |
| **Other information** | | | | | |

Intercept = Population. SD = Standard Deviation; β = estimated coefficients for the fixed effect terms; SE = Standard Error. GLMM: Generalized Linear Mixed Model; Significance codes: *** 0.001; ** 0.01; * 0.05.

**Supplementary Table S10:** Summary of the selected model for analysis of the percentage of PV cells co-expressing Fos in nucleus accumbens shell.

| | | | | | |
|---|---|---|---|---|---|
| **Method** | Generalized Linear Mixed Model (GLMM) fit by maximum likelihood | | | | |
| **Family distribution with link function** | Beta ( logit ) | | | | |
| **Formula** | response_var ~ Abstinence * Environment + (1 \| Batch) | | | | |
| **Random effects** | | **Variance** | **SD** | | |
| | Cohort (intercept) | 7.252e-09 | 8.516e-05 | | |
| **# Observations** | 41 | | | | |
| **# Cohort** | 4 | | | | |
| **Fixed effects** | | **β** | **SE** | **t value** | **Pr(>\|z\|)** |
| | Intercept | -2.2042 | 0.4154 | -5.306 | 1.12e-07 *** |
| | Abs30 | 1.8949 | 0.5216 | 3.633 | 0.00028 *** |
| | Home Cage | -1.1534 | 0.4697 | -2.455 | 0.01407 * |
| | Abs30:Home Cage | -2.0382 | 0.6757 | -3.016 | 0.00256 ** |
| **Diagnosis** | No diagnosis problem detected. | | | | |
| **Other information** | | | | | |

Intercept = Population. SD = Standard Deviation; β = estimated coefficients for the fixed effect terms; SE = Standard Error. GLMM: Generalized Linear Mixed Model; Significance codes: *** 0.001; ** 0.01; * 0.05.
